# Inactivation of p21-Activated Kinase 2 (Pak2) Inhibits the Development of *Nf2*-Deficient Malignant Mesothelioma

**DOI:** 10.1101/2020.06.30.181453

**Authors:** Eleonora Sementino, Yuwaraj Kadariya, Mitchell Cheung, Craig W. Menges, Yinfei Tan, Anna-Mariya Kukuyan, Ujjawal Shrestha, Sofiia Karchugina, Kathy Q. Cai, Suraj Peri, James S. Duncan, Jonathan Chernoff, Joseph R. Testa

## Abstract

Malignant mesotheliomas (MM) show frequent somatic loss of the *NF2* tumor suppressor gene. The *NF2* product, Merlin, is implicated in several tumor-related pathways, including p21-activated kinase (PAK) signaling. Merlin is both a phosphorylation target for PAK and a negative regulator of this oncogenic kinase. Merlin loss results in PAK activation, and PAK inhibitors hold promise for the treatment of *NF2*-deficient tumors. To test this possibility in an *in vivo* genetic system, *Nf2^f/f^;Cdkn2a^f/f^* mice were crossed to mice with conditional knockout of *Pak2*, a highly expressed group I Pak member. Cohorts of these animals were injected in either the thoracic or peritoneal cavities with adeno-Cre virus to delete floxed alleles in the mesothelial lining. Loss of *Pak2* resulted in a markedly decreased incidence and delayed onset and progression of pleural and peritoneal MMs in *Nf2;Cdkn2a-deficient (NC)* mice, as documented by Kaplan-Meier survival curves and *in vivo* bioluminescent imaging. RNA-seq revealed that MMs from *NC;Pak2^-/-^* mice showed downregulated expression of genes involved in several oncogenic pathways (Wnt, Akt) when compared to MMs from mice retaining Pak2. Kinome profiling showed that, as compared to *NC* MM cells, *NC;Pak2^-/-^* MM cells had multiple kinase changes indicative of an epithelial to mesenchymal transition. Collectively, these findings suggest that *NC;Pak2^-/-^* MMs adapt by reprogramming their kinome and gene signature profiles to bypass the need for PAK activity via the activation of other compensatory oncogenic kinase pathways. The identification of such secondary pathways offers opportunities for rational combination therapies to circumvent resistance to anti-PAK drugs.

## Introduction

Malignant mesotheliomas (MMs) are medically unresponsive cancers of the membranes lining the serous cavities. MMs often occur after chronic exposure of mesothelial cells to asbestos fibers (1). MM causes more than 3,000 deaths annually in the U.S., and a significant increase in MM incidence is predicted in certain developing countries where asbestos usage is increasing at an alarming rate and where protection of workers is minimal.

Genetically, MM is characterized by frequent somatic loss/inactivation of certain tumor suppressor genes, prominent among them being *BAP1, NF2,* and *CDKN2A/B* (2–10). *NF2* mutations and loss of Merlin expression have been reported in up to ~55% of MM cell lines (5). Among pleural MM tumors characterized by The Cancer Genome Atlas (TCGA), monoallelic deletions of *NF2* were observed in 34% of samples and biallelic inactivation in another 40% of tumors, with many of the latter harboring mutations of one allele (10). Underscoring the relevance of *NF2* inactivation to MM pathogenesis, heterozygous *Nf2* knockout mice treated with asbestos develop MM at a significantly higher frequency and markedly accelerated rate than their wild-type counterparts (6,11). Moreover, in one of these studies, 9 of 9 MM cell lines established from neoplastic ascites of *Nf2^+/-^* mice exhibited loss of the wild-type *Nf2* allele, and expression of the *Nf2* protein product, Merlin, was absent in these cells (6).

Merlin has been implicated in various tumor-related signaling pathways, prominent among them being p21-activated kinase (Pak) and Hippo signaling. Merlin regulates the protein kinases Mst1 and Mst2 (mammalian sterile 20-like 1 and −2; *a.k.a.* serine/threonine protein kinase Stk4 and Stk3) and the serine/threonine kinases Lats1 and −2 (large tumor suppressor 1 and −2). Merlin and each of these kinases are components of the highly conserved Hippo signaling pathway, which regulates organ size in *Drosophila* and mammals. Combined Mst1/2 deficiency in the liver results in loss of inhibitory phosphorylation of the downstream oncoprotein Yap1 and development of hepatocellular carcinoma (12). *Nf2*-deficient phenotypes in multiple tissues were suppressed by heterozygous deletion of *Yap1,* suggesting that Yap is a major effector of Merlin in growth regulation. The Hippo tumor-suppressive signaling pathway has also been connected with Merlin-deficient MM (13,14). Other work has shown that Merlin suppresses tumorigenesis by activating upstream components of the Hippo pathway by inhibiting the E3 ubiquitin ligase CRL4(DCAF1) (15).

Merlin is regulated by phosphorylation, with hypophosphorylated Merlin being the growth-inhibitory, functionally active tumor suppressor form, whereas hyperphosphorylated Merlin is growth-permissive (16,17). PAK directly phosphorylates Merlin at serine residue 518, a site that regulates Merlin activity and localization (18,19). Such phosphorylation of Merlin weakens its head-to-tail self-association and its association with the cytoskeleton (20). Furthermore, functional analysis of serine 518 phosphorylation has demonstrated that expression of a phospho-mimic mutant (Merlin S518D) caused striking changes in cell proliferation and shape, stimulating the creation of filopodia (21). These results strongly suggest that Merlin’s growth and motility suppression functions are attenuated following phosphorylation by Pak. Moreover, it is noteworthy that many MM tumors that lack *NF2* mutations nevertheless show functional inactivation of Merlin via constitutive phosphorylation of Ser518, in some cases as a result of inhibition of a Merlin phosphatase (22).

PAKs are serine/threonine protein kinases that are binding partners for the small GTPases Cdc42 and Rac, and they represent one of the most highly conserved effector proteins for these enzymes (23). Mammalian tissues contain six Pak isoforms: group I (Pak1, −2, and −3) and group 2 (Pak4, −5, and −6). These two groups differ substantially in form and function, and only group I Paks appear to be involved in Merlin signaling (24). Group I Paks, in particular Pak1, have oncogenic properties when expressed at high levels. In most settings, expression of Pak1 stimulates cell proliferation, survival, and motility (24,25). Group I Paks are frequently activated in human MM tumors, and genetic or pharmacologic inhibition of Paks is sufficient to inhibit MM cell proliferation and survival (26). Importantly, Merlin is more than just a target for Pak; it is also a negative regulator of this kinase. Merlin binds to and inactivates Pak1, and loss of Merlin results in the activation of Pak (20). These results suggest that the growth and motility abnormalities of *Nf2*-null cells might in part be attributed to the activation of Pak and its downstream targets. These data also suggest that inhibitors of Pak might be useful in switching off such signaling pathways. We hypothesized that loss of Pak activity would counteract some aspects of Nf2/Merlin loss-of-function by compromising key downstream oncogenic signaling pathways in mesothelial cells. To test this possibility, we crossed *Nf2^f/f^;Cdkn2a^f/f^* mice to mice with conditional deletion of *Pak2,* which encodes the most highly expressed group I Pak isoform in most tissues. Cohorts of these animals were then injected in either the thoracic or peritoneal cavities with adeno-Cre virus to delete floxed alleles in the mesothelial lining, with the goal being to determine if Pak2 loss diminishes MM onset and/or progression. We show that loss of Pak2 indeed delays MM tumorigenesis, though studies employing multiplexed kinase inhibitor beads and mass spectrometry (MIB/MS) and RNA-seq technologies revealed that MM cells ultimately reprogram their kinome and gene signature profiles to bypass the need for Pak2 activity. The identification of such secondary pathways offers opportunities for the rational design of combination therapies to circumvent resistance to anti-Pak drugs.

## Materials and Methods

### Mouse strains

*LucR;Nf2^f/f^;Cdkna^f/f^* mice in FVB/N genetic background (27), a kind gift of Dr. Anton Berns, were maintained in our laboratory in a mixed FVB/N × 129/Sv background. The floxed *Cdkn2a* locus permits excision of exon 2, resulting in inactivation of both p16(Ink4a) and p19(Arf) tumor suppressors. *Pak2^f/f^* mice in a C57BL/6J background (28) were crossed a minimum of five generations to *Nf2^f/f^;Cdkna^f/f^* mice in a FVB/N × 129/Sv background. All mouse studies were performed in accordance with protocol #18-03 approved by the Fox Chase Cancer Center (FCCC) Institutional Animal Care and Use Committee (IACUC).

### Adeno-Cre injections

For studies involving intrathoracic (IT) injections of adeno-Cre virus, animals with *Nf2^f/f^;Cdkna^f/f^;Pak2^+/+^* and *Nf2^f/f^;Cdkna^f/f^;Pak2^f/f^* genotypes were divided into two cohorts with equivalent numbers of male and female mice in each group. For intraperitoneal (i.p.) injection studies, animals with *Nf2^f/f^;Cdkna^f/f^* and *Nf2^f/f^;Cdkna^f/f^;Pak2^f/f^* genotypes were again divided into two cohorts as above. Ad5CMVCre (adeno-Cre) virus was purchased from the Viral Vector Core of the University of Iowa. At 8-10 weeks of age, all mice were injected either IT or i.p. with adeno-Cre virus (50 μl of 3-6 × 10^10^ PFU) in PBS, with approximately equal numbers of mice of each gender in each cohort. Expression of adeno-Cre results in the removal of the floxed exons in the *Nf2, Cdkn2a,* and *Pak2* loci, resulting in the following genotypes in adeno-Cre-infected cells: *Nf2^Δ/Δ^;Cdkn2a^Δ/Δ^* and *Nf2^Δ/Δ^;Cdkn2a^Δ/Δ^;Pak2^Δ/Δ^*, and the corresponding mice are referred to as *NC;P^wt^* and *NC;P^-/-^* mice hereafter). All mice were closely monitored for tumor formation over a period of up to 12 months. Mice were monitored daily and were immediately sacrificed by CO_2_ inhalation upon signs of pain/distress or illness as judged by lethargic behavior, weight loss or bloating, difficulty in breathing, hunched posture, rough hair coat, dehydration or detectable tumor volume approaching 10% of overall body weight. Tissues of all organs of the pleural and peritoneal cavities were collected from sacrificed mice, and tumor specimens were subjected to histopathological assessment by an experienced animal pathologist (K.Q.C.) of FCCC’s Histopathology Facility, a core service supported by our NCI Cancer Center Support Grant. Portions of tumors were also saved in both O.C.T. Compound and RNAlater Solution (Thermo Fisher, Waltham, MA) and immediately frozen at −80°C for subsequent study. When possible, portion of the tumor was also disaggregated and cultured to generate tumor cell lines.

### Detection of tumors and statistical considerations

Survival curves for *NC;P^wt^* versus *NC;P^-/-^* mice were compared using one-sided, log-rank tests. Kaplan-Meier plots were used to display time to tumor detection among the two separate cohorts for both IT and i.p. injection studies.

### Tumor histopathology, immunohistochemistry and RT-PCR

Tumor tissues were paraffin embedded, sectioned and deparaffinized, followed by staining of sections with hematoxylin and eosin (H&E) for histopathologic evaluation. Other sections were for immunohistochemistry (IHC), performed using standard methods. To confirm the diagnosis of MM, IHC was performed for various MM markers, including mesothelin, detected with D-16 antibody (Santa Cruz Biotechnology, Dallas, TX), and cytokeratin 8, detected with TROMA-1 antibody (DSHB, University of Iowa, Iowa City, IA). To evaluate tumor cell proliferation, IHC staining was performed with antibodies for Ki-67 (Dako/Agilent, Santa Clara, CA). In some tumors, reverse transcription-PCR (RT-PCR) analysis was also performed to confirm the diagnosis of MM, using primers for the MM markers Wt1 and Msln (mesothelin). Specific primers used for RT-PCR are shown in Supplemental Table S1.

### Preparation of luciferin and in vivo bioluminescent Imaging (BLI)

*LucR;Nf2^f/f^;Cdkna^f/f^* mice were crossed to *Pak2^f/f^* mice to generate offspring with different *Pak2* genotypes (^+/+^, ^+/*f*^,^*f/f*^), which at 8-10 weeks of age were injected IT with adeno-Cre virus. In addition to excising the floxed *Nf2*, *Cdkna,* and *Pak2* alleles in mesothelial cells lining the pleural cavity, expression of Cre recombinase also removes a floxed polyadenylation acid sequence before the ORF of the luciferase reporter transgene *(LucR),* thereby permitting luciferase expression for monitoring tumor progression. Beginning at 6-7 weeks after injecting with adeno-Cre virus, littermates with different genotypes were injected i.p. with 150 mg of filtered D-Luciferin, Firefly, potassium salt (Caliper Life Sciences) in PBS per kg mouse body weight 10 minutes before imaging. BLI scans were acquired using an IVIS Spectrum Imaging System (Caliper Life Sciences) as described (29) to assess the presence and relative size of tumors, as indicated by the intensity of luminescent signals detected. The same mice were imaged weekly until tumors began to form. The experiment was repeated four times.

### Preparation of lentivirus expressing shRNA against *Pak2*

A Tet-pLKO-puro plasmid was a gift from Dmitri Wiederschain (Addgene, plasmid # 21915). The shRNAs targeting mouse *Pak2* were created by the cloning of annealed forward and reverse oligos synthesized based on information provided at the Broad Institute’s website (https://portals.broadinstitute.org/gpp/public/). The clone numbers and target sequences are as follows: mouse shPak2 #70 (TRCN0000432870): TTCGGATGAGCAGTACCATTT; mouse shPak2 #85 (TRCN0000417285): ATGATTGATGTAGCTCTTTAC. The shRNAs was cloned as previously described (30). Lentivirus was produced by transfecting 293T cells with the two different Tet-inducible shPak2, or control shGFP, and the packaging plasmids pMD2G and pPax2, using Lipofectamine 2000 transfection reagent (11668019; Thermo Fisher). After 24 h and 48 h, virus particles were collected, filtered through a 0.45-μm PES filter, and then used to infect cell lines for experiments described below.

### *In vivo* model to assess lung tumor burden of MM cells following knock down of *Pak2*

In this experiment, we used asbestos-induced mouse MM cells (MM87) from *Nf2^+/-^;Cdkn2a^+/-^* mice (31). The MM87 cells were infected with four separate tet-inducible lentiviruses against *Pak2* to assess knockdown of *Pak2.* The clones with the most robust knockdown of *Pak2* were expanded, and each clone was injected into the tail vein of three NSG mice (0.5 x 10^6^ cells/mouse), followed by i.p. administration of doxycycline beginning after 7 days and then every 2 days thereafter. Animals were sacrificed on day 21, and then the lungs were harvested for histopathological assessment of tumor colonization. An unpaired t-test with Welch’s correction was used to determine the statistical significance of the data.

### Immunoblot analysis and antibodies

For immunoblotting, protein lysates were prepared as previously described (31). Lysates (30-50 μg/sample) were loaded on gels, transferred to nitrocellulose membranes (1620115, Bio-Rad, Hercules, CA), and probed with anti-Nf2 (D1D8, 6995, 1:4000), anti-Pak2 (2608, 1:2000), anti-Akt (9272, 1:2000), anti-phospho (p)-Akt (Ser473) (B9E)XP (4060, 1:4000), anti-p44/42 MAPK (Erk1-2) (9102, 1:2000), Pdgfra (D1E1E) (XP 3174, 1:2000), Pdgfrβ (C82A3) (4564, 1:2000), anti-Limk1 (3842, 1:1000), anti-MKK6 (9264, 1:1000), anti Ddr1 (D1G6) (XP 5583, 1:1000), E-Cadherin (24E10, 3195, 1:1000), anti-Slug (C19G7, 9585, 1:1000), anti-Snail (C15D3, 3879, 1:1000), anti-Stat3 (124H6, 9139, 1:5000), anti-p-Stat3 (Tyr705) (D3A7, XP 9145, 1:1000), anti-Fyn (4023, 1:1000), anti-Met (25H2, 3127, 1:1000), anti-Jak1 (6G4, 3344, 1:1000), and anti-N-Cadherin (D4R1H, XP 13116, 1:1000) from Cell Signaling Technology (Danvers, MA); anti-phospho-Pak1-2-3 (pSer141) (44-940G, 1:2000) from Invitrogen/Thermo Fisher Scientific (Carlsbad, CA); recombinant anti-p16Ink4a (ab211542, 1:1000) from Abcam (Cambridge, MA); anti-p19Arf (5-C3-1) (sc-32748, 1:500), anti-p-Erk (E-4) (sc-7383, 1:2000), anti-Gapdh (6C5, sc-32233, 1:50,000), anti-β-catenin (E-5) (sc-7963, 1:1000) and anti-β-actin (C4, sc-47778, 1:50,000) from Santa Cruz Biotechnology, and anti-DDR2, Clone 2B12.1, MABT322, 1:1000. Anti-Rabbit IgG, peroxidase-linked species-specific whole antibody (from donkey), secondary antibody (45-000-682, 1:5000) and anti-Mouse IgG, peroxidase-linked species-specific whole antibody (from sheep) secondary antibody (45-000-679, 1:5000-50000 depending on the primary antibody) were both from Fisher Scientific (Waltham, MA). Immunoblots were imaged using Immobilon Western Chemiluminescent HRP Substrate (ECL) (WBKLS0500, MilliporeSigma, Ontario Canada).

### Migration assay

*In vitro* migration of MM cells from *NC;P^wt^* mice and *NC;P^-/-^* mice was measured using a Transwell Multiwell Plate with Polyester Membrane Inserts (Corning, Edison, NJ). In the upper compartment of each well, 8.0 × 10^3^ cells/well were seeded in serum-free DMEM medium, and in the lower compartment we placed DMEM medium containing 10% serum. Cells were incubated at 37°C in a humidified 5% CO_2_ incubator for 22 hours, and then the permeable insert was stained according to the manufacturer’s recommendations. The migrating cells on the underside of the membrane were fixed and stained with Diff-Quik solution.

### MTS assay

Lentivirus particles harboring two different Tet-inducible shPak2 constructs (#70, #85), or control shGFP, were used to infect pleural *NC;P^wt^* MM cell line #2. The cells were seeded on a 6-well plate and allowed to proliferate to approximately 50% confluency. Then virus particles and polybrene (sc-134220; Santa Cruz) at a concentration of 2 μg/ml were added to the plate, which was then spun down for 2 h at 2000 rpm and incubated for 24 h at 37°C in a humidified 5% CO_2_ incubator. The cells were then selected in fresh media containing puromycin (4μg/ml) for 3 days. When the selection was completed, the cells were treated with doxycycline for 24 h and seeded in 96-well plates at 500 cells/well. The following day, the cells were refed with fresh media containing puromycin (4 μg/ml) and doxycycline (Sigma, D9891; stock 1 mg/ml, diluted 1:1000), which was also added to the cells daily thereafter. Cell viability was assessed at 2, 4, 5 and 6 d using a CellTiter 96^®^ AQueous One Solution Cell Proliferation Assay (MTS) (G3582; Promega, Madison, WI). Cells were incubated with the MTS reagent 3-4 h, and the OD value at 490 nm was measured, using a 96-well microplate reader (BioRad, Santa Monica, CA).

### RNA-seq and gene set enrichment analyses

Tumor cell RNA was isolated in Trizol and purified using RNeasy columns (Qiagen, Germantown, MD). An initial RNA-seq analysis was performed on a small set of MMs, with mRNA-seq libraries prepared as previously described (32), followed by loading onto an Illumina NextSeq 500 sequencer. RNA-seq analysis was performed on an expanded set of MMs by Novogene (Sacramento, CA), using its Illumina NovaSeq 6000 platform. The Fastq files were aligned to the mm10 mouse genome using the STAR RNA-seq aligner. Raw sequence counts for each gene were produced with HTseq (https://htseq.readthedocs.io), and differentially expressed genes were determined using DESeq2 (33) For functional enrichment analysis, genes identified as differentially expressed with nominal p-value < 0.5 were ranked by fold-change and mapped to human genes using R biomaRt (34). Next, these genes were analyzed using the GSEAPreranked method of Gene Set Enrichment Analysis (GSEA) (35), with “classic” enrichment statistic, applied to the curated canonical pathways (c2cp) and Gene Ontology gene sets from the Molecular Signature Database (MSigDB). Heatmaps were made using pheatmap library available through Bioconductor (https://www.bioconductor.org). Raw sequencing data is in the process of being deposited in the GEO repository.

### MIBs preparation and chromatography

In an attempt to gain an unbiased and more comprehensive view of Merlin signaling, kinome reprogramming analysis was performed using MIB/MS as previously described (36). In brief, cells were lysed on ice in buffer containing 50 mM HEPES (pH 7.5), 0.5% Triton X-100, 150 mM NaCl, 1 mM EDTA, 1 mM EGTA, 10 mM sodium fluoride, 2.5 mM sodium orthovanadate, 1X protease inhibitor cocktail (Roche), and 1% each of phosphatase inhibitor cocktails 2 and 3 (Sigma). Particulate was removed by centrifugation of lysates at 21,000 g for 15 min at 4°C and filtration through 0.45 μm syringe filters. Protein concentrations were determined by BCA analysis (Thermo Scientific). Endogenous kinases were isolated by flowing lysates over kinase inhibitor-conjugated Sepharose beads (purvalanol B, VI16832, PP58 and CTx-0294885 beads) in 10 ml gravity-flow columns. After 2×10-ml column washes in high-salt buffer and 1×10-ml wash in low-salt buffer (containing 50 mM HEPES (pH 7.5), 0.5% Triton X-100, 1 mM EDTA, 1 mM EGTA, 10 mM sodium fluoride, and 1M NaCl or 150 mM NaCl, respectively, retained kinases were eluted from the column by boiling in 2 x 500 μl of 0.5% SDS, 0.1 M TrisHCl (pH 6.8), and 1% 2-mercaptoethanol. Eluted peptides were reduced by incubation with 5 mM DTT at 65°C for 25 min, alkylated with 20 mM iodoacetamide at room temperature for 30 min in the dark, and alkylation was quenched with DTT for 10 min. Samples were concentrated to approximately 100 μl with Millipore 10kD cutoff spin concentrators. Detergent was removed by chloroform/methanol extraction, and the protein pellet was resuspended in 50 mM ammonium bicarbonate and digested with sequencing-grade modified trypsin (Promega) overnight at 37°C. Peptides were cleaned with PepClean C18 spin columns (Thermo Fisher Scientific), dried in a speed-vac, resuspended in 50 μl of 0.1% formic acid, and extracted with ethyl acetate (10:1 ethyl acetate:H_2_O). Briefly, 1 mL ethyl acetate was added to each sample, vortexed, and centrifuged at maximum speed for 5 min, and then removed. This process was repeated 4 more times. After removal of ethyl acetate following the fifth centrifugation, samples were placed at 60°C for 10 min to evaporate residual ethyl acetate. The peptides were then dried in a speed vac, and subsequent LC-MS/MS analysis was performed.

### Nano-LC-MS/MS

Proteolytic peptides were resuspended in 0.1% formic acid and separated with a Thermo Scientific RSLC Ultimate 3000 on a Thermo Scientific Easy-Spray C18 PepMap 75 μm x 50 cm C-18 2 μm column with a 240 min run time on a gradient of 4-25% acetonitrile with 0.1% formic acid at 300 nL/min at 50°C. Eluted peptides were analyzed by a Thermo Scientific Q Exactive plus mass spectrometer utilizing a top 15 methodology in which the 15 most intense peptide precursor ions were subjected to fragmentation. The AGC for MS1 was set to 3×10^6^ with a maximum injection time of 120 ms, the AGC for MS2 ions was set to 1×10^5^ with a maximum injection time of 150 ms, and the dynamic exclusion was set to 90 s.

### Data processing for kinome profiling analysis

MIB/MS was performed in biological triplicates for each condition and analyzed by LC-MS/MS in technical duplicates. Raw data analysis of MIB/MS experiments was performed using MaxQuant software 1.6.1.0 and searched using Andromeda 1.5.6.0 against the Swiss-Prot human protein database (downloaded on July 26, 2018). The search was set up for full tryptic peptides with a maximum of two missed cleavage sites. All settings were default and searched using acetylation of protein N-terminus and oxidized methionine as variable modifications. Carbamidomethylation of cysteine was set as fixed modification. The precursor mass tolerance threshold was set at 10 ppm and maximum fragment mass error was 0.02 Da. Label-free quantification was performed using MaxQuant. The match between runs was employed, and the significance threshold of the ion score was calculated based on a false discovery rate of < 1%.

For MIB/MS data analysis, MaxQuant normalized LFQ values were imported into Perseus software (1.6.2.3) for quantitation. MIB/MS profiles were processed in Perseus software in the following manner: normalized MIB/MS LFQ ratios were log2 transformed, MIB/MS technical replicates averaged, rows filtered for minimum valid kinases measured (n=>3 kinases) and normalized by Z-score. Principal component analysis and hierarchical clustering (Euclidean) of kinase log2 LFQ z-scores was then performed to visualize kinome profiles amongst samples. Differences in kinase abundance among sample conditions were determined using a two-sample Student’s t-test with the following parameters, (S0 0.1, and Side, Both) using Benjamini-Hochberg FDR 0.05 using Perseus software. The mass spectrometry proteomics files are currently in the process of being deposited to the ProteomeXchange Consortium via the PRIDE partner repository.

## Results

### Inactivation of Pak2 results in decreased incidence and delayed onset and progression of pleural and peritoneal MMs in Nf2-deficient mice

Locotemporal expression of Cre recombinase by either IT or i.p. injection of adeno-Cre virus was used to induce mesothelial cell-specific homozygous deletions of *Nf2*, *Cdkn2a,* with or without excision of *Pak2,* in homozygous compound CKO mice. Examples of genotyping of *Nf2^f/f^;Cdkna^f/f^;Pak2^+/+^* and *Nf2^f/f^;Cdkna^f/f^;Pak2^f/f^* mice and immunoblotting demonstrating loss of expression of conditionally knocked out genes in MMs arising after injection of adeno-Cre virus in these animals are depicted in Fig. 1A and 1B, respectively. The IT-injected mice were followed for up to one year, and by the end of the study, 16 of 19 *NC;P^wt^* animals (84%) developed pleural MM, with a median survival of 26 weeks. In contrast, only 8 of 15 (53%) *NC;P^-/-^* mice developed pleural MM, and the median survival was prolonged to 34 weeks. Among animals injected i.p. with adeno-Cre virus, 18 of 22 (82%) *NC;P^wt^* mice developed peritoneal MM (median survival: 24 weeks) versus only 10 of 23 (43%) *NC;P^-/-^* mice (median survival: 35 weeks). Kaplan-Meier survival curves of IT- and i.p.-injected mice that developed MM are shown in Figure 2A and 2B, respectively. In both the IT and the i.p. studies, the incidence of MM was much lower in *NC;P^-/-^* mice than in the *NC;P^wt^* cohort. Moreover, statistical differences in the survival of *NC;P^wt^* versus *NC;P^-/-^* mice with MM were highly significant in both IT and i.p. studies. MMs from *NC;P^wt^* and *NC;P^-/-^* mice showed similar histopathology, with the vast majority of tumors being sarcomatoid (Supplemental Figure S1), which is consistent with our previous studies of MMs in conditional NC mice (32).

**Figure 1.**
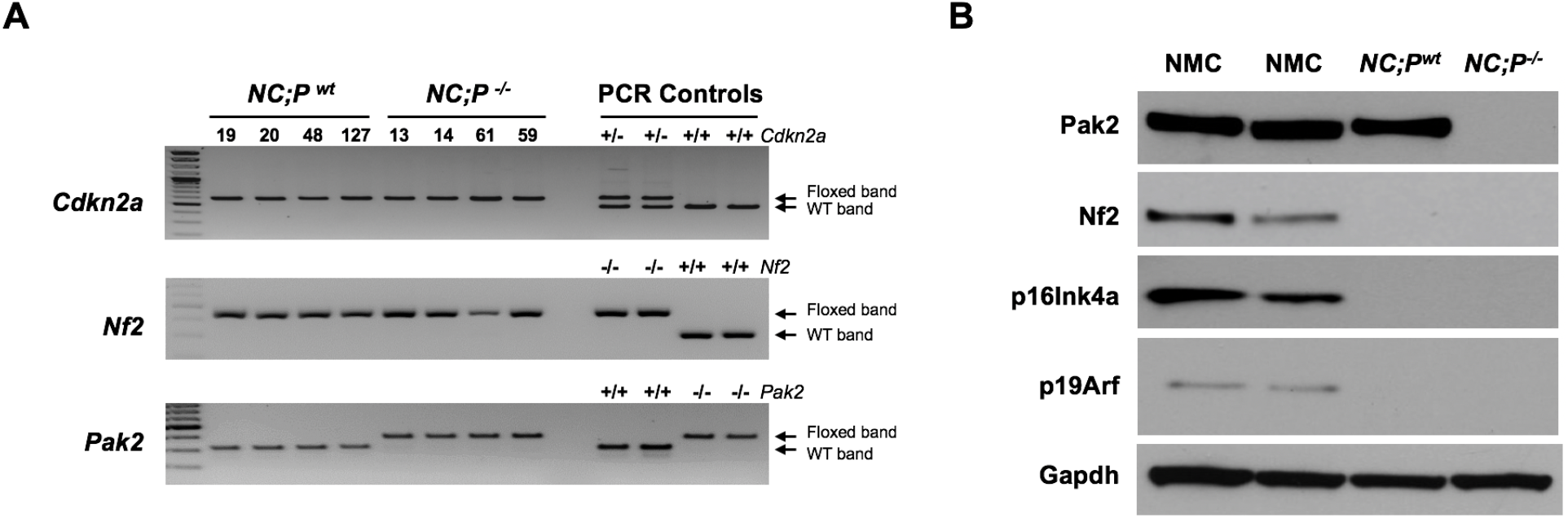
Genotyping of *Nf2^f/f^;Cdkna^f/f^;Pak2^+/+^* and *Nf2^f/f^;Cdkna^f/f^;Pak2^f/f^* mice injected intrathoracically (IT) or intraperitoneally (i.p.) with adeno-Cre virus and immunoblotting demonstrating loss of expression of conditionally knocked out genes in malignant mesotheliomas (MMs) from these mice. (**A**) Genotyping of tail DNA from four representative *Nf2^f/f^;Cdkna^f/f^;Pak2^+/+^* mice and four *Nf2^f/f^;Cdkna^f/f^;Pak2^f/f^* mice that developed peritoneal MM after i.p. injection of adeno-Cre virus. PCR controls for floxed and wild type alleles of *Cdkn2a, Nf2* and *Pak2* were from tail DNA of control heterozygous and homozygous mice conditional knockout mice. (**B**) Immunoblotting of early passage cell lines derived from peritoneal MMs of *NC;P^-/-^* and *NC;P^wt^* mice. NMC, normal (mouse) mesothelial cells.

**Figure 2.**
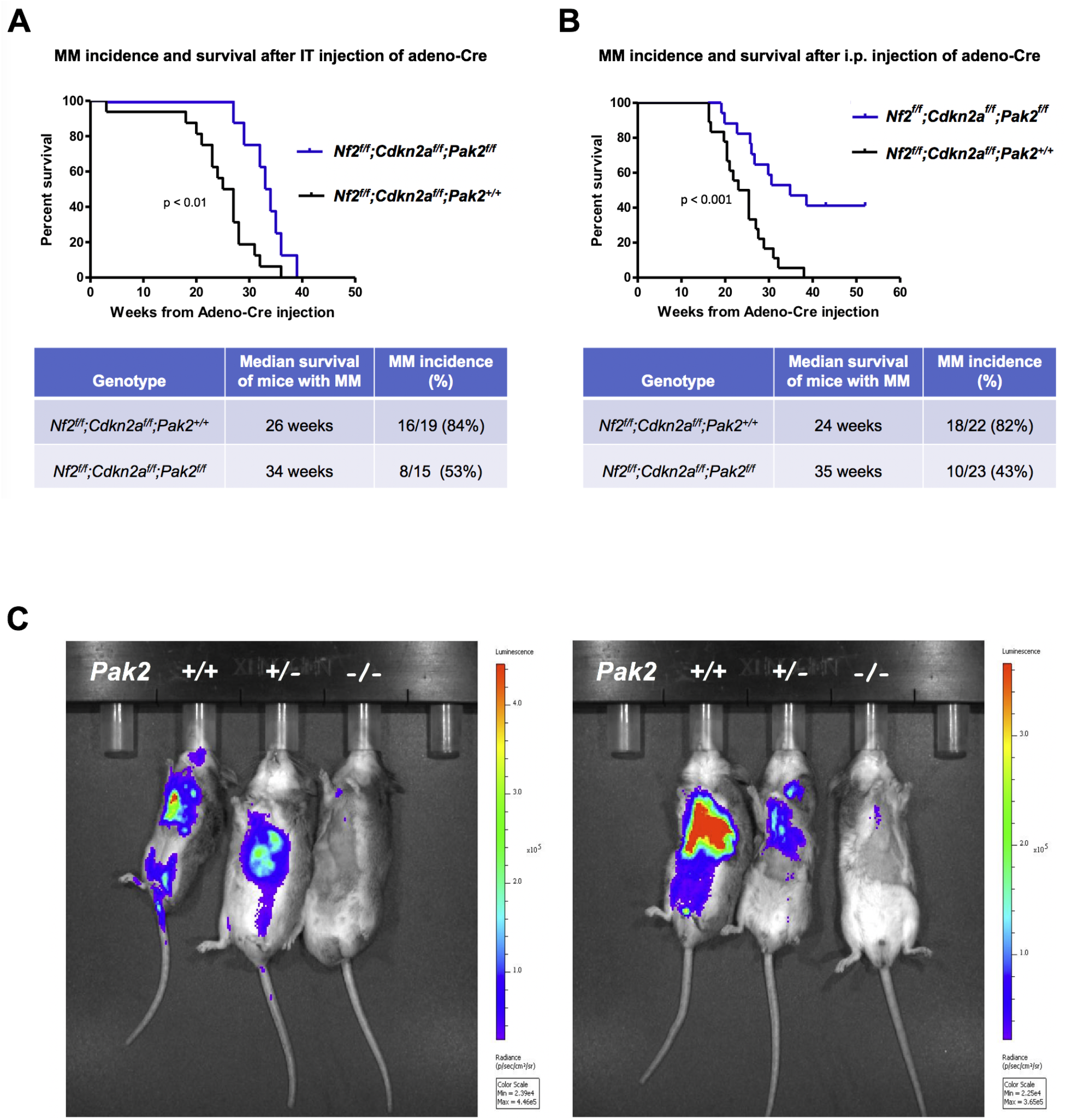
MM progression, incidence and Kaplan-Meier survival curves of cohorts of conditional *Nf2^f/f^;Cdkna^f/f^;Pak2^+/+^* and *Nf2^f/f^;Cdkna^f/f^;Pak2^f/f^* mice injected intrathoracically (IT) or intraperitoneally (i.p.) with adeno-Cre virus. (**A**) MM incidence and survival of mice injected IT with virus and succumbing to MM. (**B**) MM incidence and survival in mice injected i.p. The difference in the incidence of MM between *Pak2^wt^* and *Pak2^-/-^* mice injected i.p. was highly significant (p-value < 0.01), while the p-value for the difference in the incidence of MM between Pak2^wt^ and Pak2^-/-^ mice injected IT was <0.06 (Fisher’s exact test). (**C**) Bioluminescent imaging reveals delayed MM tumor progression in *Nf2*-null mice following excision of one or both alleles of *Pak2.^ff^LucR;Nf2^f/f^;Cdkn2a^f/f^* mice were crossed to *Pak2^f/f^* mice to generate offspring having wild type (^+/+^) *Pak2* or with one or both floxed (*^+/f^* or ^*f/f*^, respectively) *Pak2* alleles. Mice were injected intrathoracically (IT) with adeno-Cre virus to excise floxed alleles of thoracic mesothelial lining cells. Infection with adeno-Cre virus also removes a floxed polyadenylation sequence before the ORF of a luciferase reporter transgene *(LucR).* The latter permits luciferase expression to monitor tumor progression, using D-luciferin as a substrate and bioluminescent imaging with an IVIS Imaging System. Shown is bioluminescent imaging on two sets of ^*ff*^*LucR;NC* littermates with three different *Pak2* genotypes. The mice were injected with D-luciferin substrate 6 months *(left* panel) or 7 months *(right)* after IT injection of adeno-Cre virus; mice with excision of Pak2 show delayed tumor progression as indicated by reduced intensity of luminescent signals.

### In vivo bioluminescent imaging confirms that loss of Pak2 delays MM progression in Nf2-deficient mice

BLI scanning revealed intense luminescent signals in *NC;P^wt^* mice beginning at about 6 months, with less signal observed in *NC;P2^+/-^* (heterozygous loss of *Pak2)* littermates (Figure 2C). BLI scanning was repeated with four sets of littermates with different *Pak2* genotypes, and in each instance tumor progression, as indicated by diminished intensity of the luminescent signals observed, was consistently delayed in mice with homozygous excision of *Pak2.*

### Knockdown of Pak2 diminishes metastatic colonization of Nf2-deficient MM cells in the lung

MM cells (MM87), which were previously derived from an asbestos exposed *Nf2^+/-^;Cdkn2a^+/-^* mouse, were infected with tet-inducible lentiviruses against *Pak2.* Two clones that demonstrated marked knockdown of *Pak2* (shPak2 85 and shPak2 70 in Figure 3A) were expanded and injected individually into the tail vein of NSG mice. The mice were injected with doxycycline i.p. after 7 days and then every 2 days thereafter; all animals were sacrificed on day 21, and lungs were examined histopathologically for tumor colonization (Figure 3B). H&E staining revealed that tumor colonization of the lungs by MM87 cells was consistently diminished in the cell clones expressing shRNA against *Pak2* versus control MM87 cells infected with lentivirus against GFP (Figure 3C), and the differences were statistically significant (p < 0.05).

**Figure 3.**
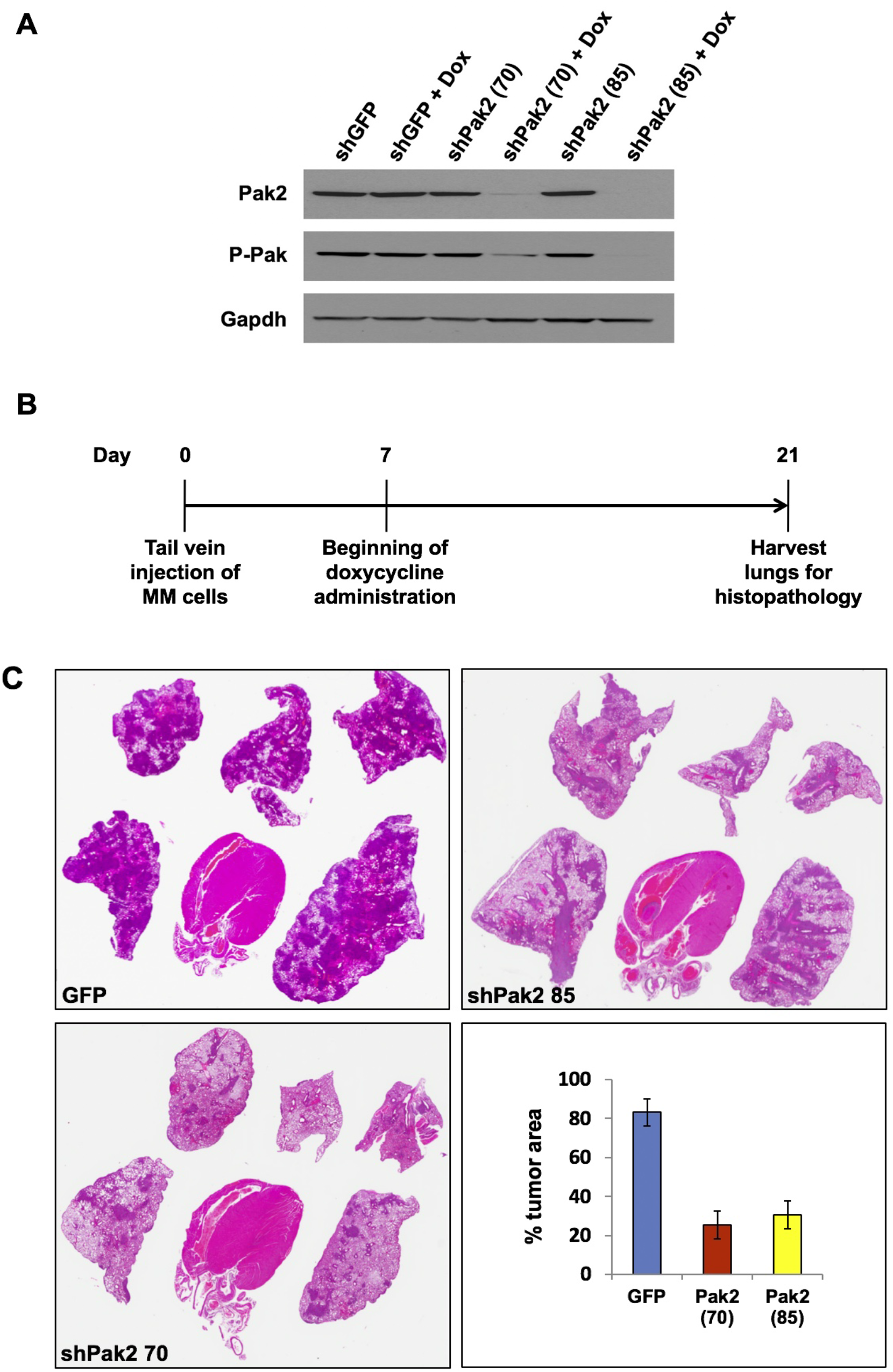
Knockdown of *Pak2* diminishes metastatic colonization of *Nf2*-deficient MM cells in the lung. Cell line MM87, which was derived from asbestos-induced MM from a *Nf2^+/-^;Cdkn2a^+/-^* mouse, were infected with tet-inducible lentiviruses against *Pak2.* (**A**) Immunoblot demonstrating expression of Pak2 after knockdown with shRNA. Two clones with robust knockdown of *Pak2* (shPak2 85 and shPak2 70) and one clone infected with lentivirus against GFP, used as a control, were selected for tail vein injections into NSG mice. (**B**) MM clones were each injected into the tail vein of three different NGS mice, followed by injection with doxycycline 7 days later and every 2 days thereafter; all animals were sacrificed on day 21, and lungs were collected for histopathological assessment of tumor burden. (**C**) H&E staining illustrating representative tumor colonization (darkly stained areas) of lungs by MM87 cells expressing shGFP, shPak2 85, or shPak2 70. Bar graph of percent of lung consisting of tumor is shown in the panel at the lower right.

### Loss of Pak2 results in decreased tumor cell migration, cell viability and Erk activity

We compared cell migration of MM cells from *NC;P^wt^* mice versus MM cells from *NC;P^-/-^* mice. Twenty-two hours after seeding, the cells were evaluated for migratory ability using a transwell migration assay, as described in the Materials and Methods. As shown in Figure 4A, the *NC;P^-/-^* MM cell line tested showed markedly less migration than a *NC;P^wt^* MM cell line. We also found that Erk activity was much lower in *NC;P^-/-^* MM cells as compared to *NC;P^wt^* MM cells (Figure 4B). Furthermore, an MTS assay performed on *NC;P^wt^* MM cells infected with a lentivirus expressing doxycycline-inducible *shPak2* revealed a sustained decrease in cell viability beginning at 4 days after starting treatment with doxycycline (Figure 4C, D).

**Figure 4.**
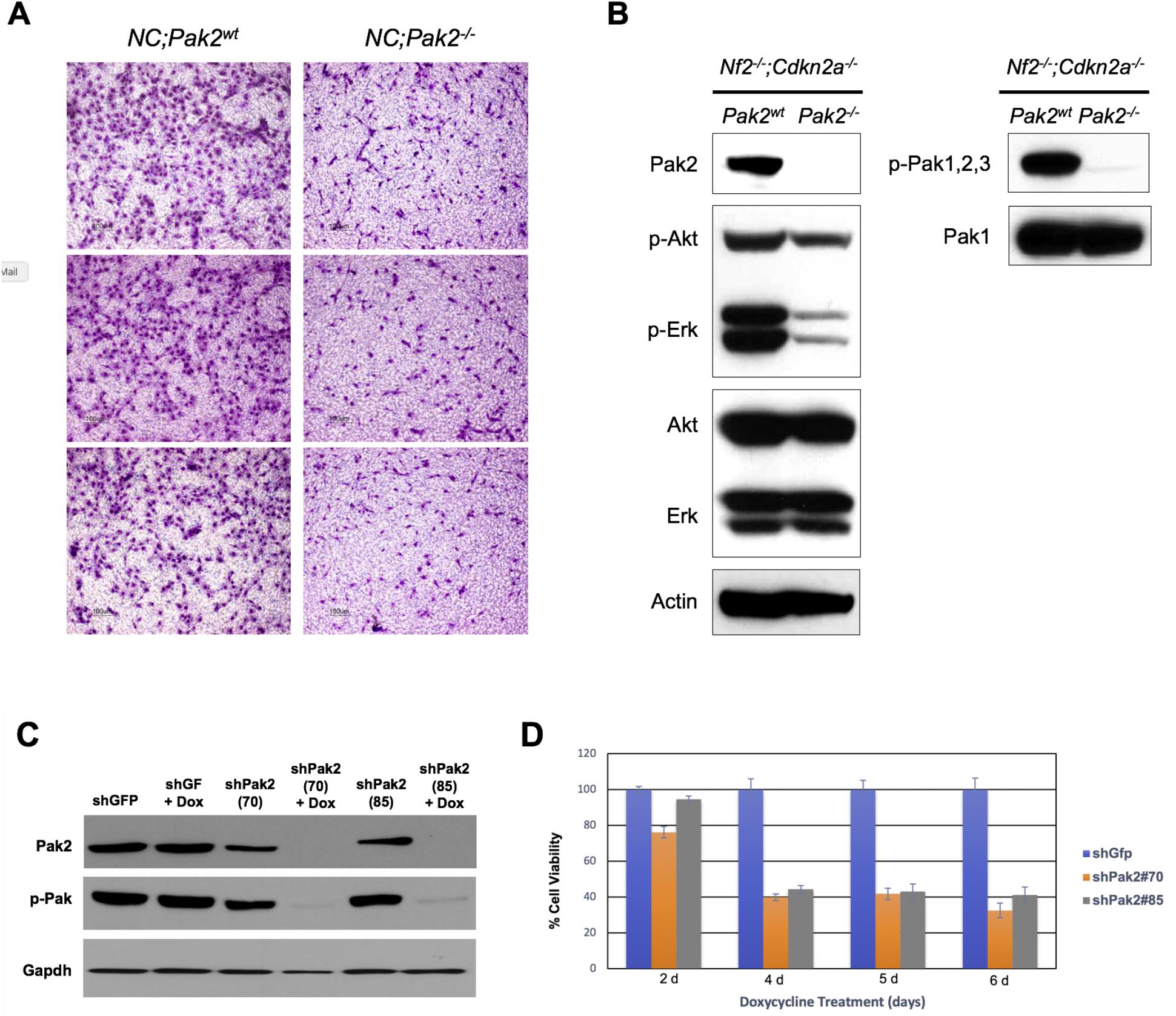
Loss of Pak2 results in decreased tumor cell migration, cell viability and Erk activity. (**A**) *In vitro* migration of MM cells from *NC;P^wt^* mice and *NC;P^-/-^* mice was measured using a transwell assay. Twenty-two hours after seeding, the MM cells were evaluated for migratory ability as described in the Materials and Methods, with cells seeded in triplicate wells. Note that *NC;P^-/-^* cells tested show markedly less migration than *NC;P^wt^* cells. (**B**) Immunoblotting illustrating decreased Erk activity in *NC;P^-/-^* MM cells than in *NC;P^wt^* MM cells. (**C**) Immunoblot demonstrating knockdown of Pak2 in *NC;P^wt^* MM cells infected with lentivirus expressing either of two doxycycline-inducible shPak2 (#70 and #85) after treatment with doxycycline for 72 h. (**D**) MTS assay performed on *NC;P^wt^* MM cells infected with lentivirus expressing doxycycline-inducible shPak2 showing sustained decrease in cell viability beginning at 4 days after starting treatment with doxycycline. Lentivirus expressing shGFP was used as a control.

### RNA-seq analysis reveals that Pak2 loss in Nf2-null MM cells results in diminished expression of genes involved in oncogenic pathways

RNA-seq analysis was performed on early passage (p 3-4) cell lines from 3 peritoneal MMs from *NC;P^wt^* mice and 5 MMs from *NC;P^-/-^* mice to identify genes that are differentially expressed due to loss of Pak2, one of the main downstream effectors of Nf2. Most of the differentially expressed genes in MMs from the *NC;P^-/-^* mice were downregulated. Numerous downregulated genes are involved in muscle contraction and cardiac epithelial-mesenchymal transition (EMT) pathways (Supplemental Figure S2), whereas many others are members of Wnt signaling *(Wif1, Nkd1, Axin2, Gli1, Gpc3,* Notum, *Sfrp5, Fzd4/8/9, Porcn, Lrp4, Wnt9b, Sost).* Other downregulated genes involved kinase and other cancer-related pathways *(Igf2, Fgf9/18, Fgfr3/4,* Notch3, *Pax7, Pik3r3, Tgfb2)* or were stem cell markers *(Sox8/11)* and integrins *(Itga7/8).* Fewer upregulated genes were observed in MMs from the *NC;P^-/-^* mice; these included a Wnt inhibitor gene (*Dkk2*), two kinase genes (Axl, Styk1), the transcription repressor gene *Foxg1,* and the dual-specificity phosphatase genes Dusp4 and Dusp5, whose products dephosphorylate MAPK proteins such as ERK. A heatmap of genes differentially expressed in MM cell lines from *NC;P^-/-^* mice versus MM lines from *NC;P^wt^* mice is shown in Figure 5, including genes involved in Wnt signaling (Figure 5A) and various other pathways (Figure 5B). Eight cancer-related genes were validated by semi-quantitative RT-PCR analysis (Figure 5C).

**Figure 5.**
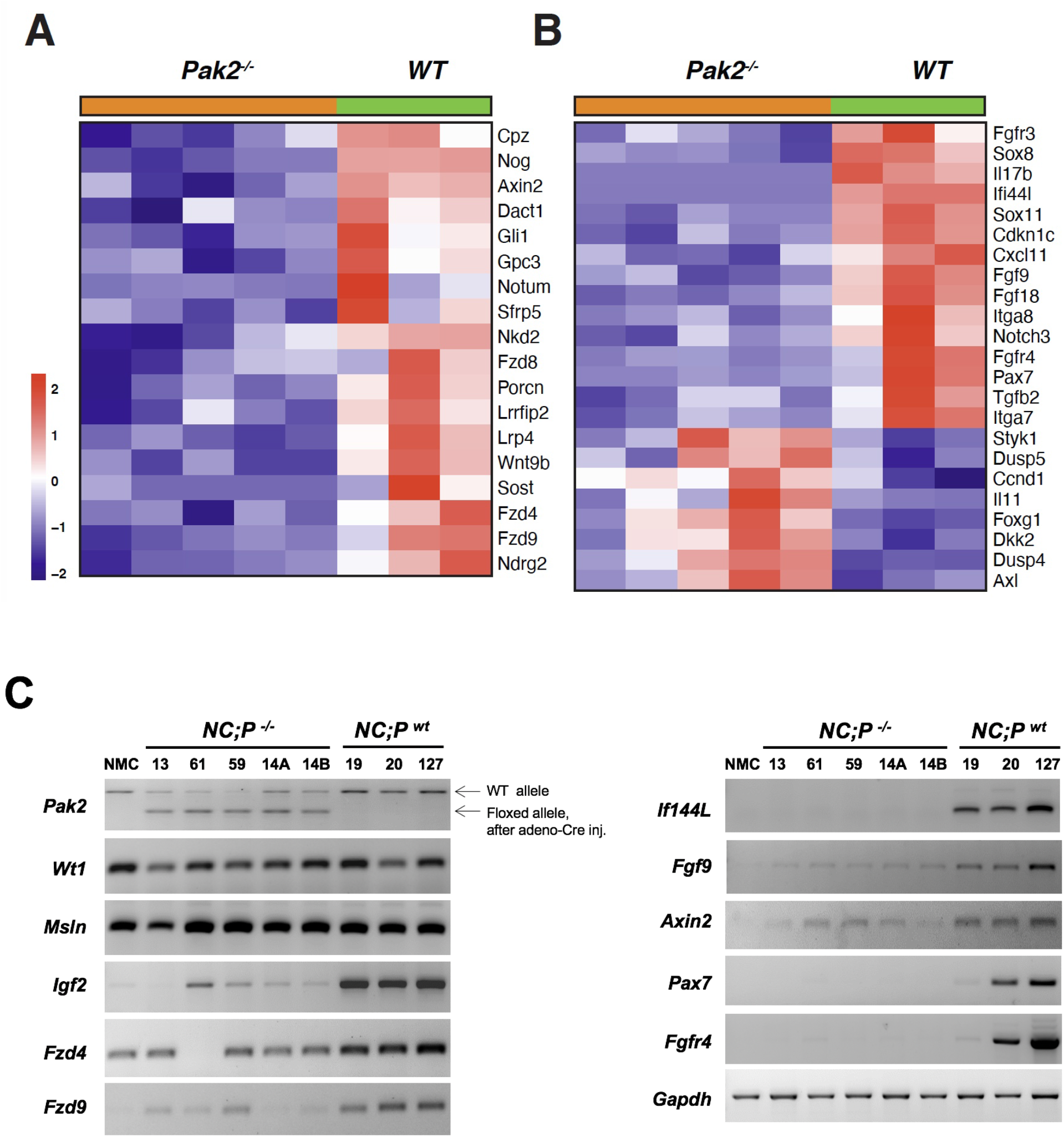
Heatmaps showing expression patterns of genes in *Pak2^-/-^* and *Pak2^wt^* tumors. Heatmaps depict differentially expressed genes observed in peritoneal MMs tumors from 3 *NC;P^wt^* mice versus 5 peritoneal MMs from *NC;P^-/-^* mice. (**A**) Genes involved in Wnt signaling pathway regulation. (**B**) Genes that involve multiple pathways including Akt and Notch signaling, cardiac EMT pathway, cell cycling, stem cell pathways, and integrins. (**C**) Validation of several differentially expressed genes by semi-quantitative RT-PCR analysis. Down-regulated genes in *NC;P^-/-^* MM cells tested include two involved in Wnt signaling, *Fzd9* and *Axin2,* several involved in Akt signaling, *Igf2, Fgf9* and *Fgfr4,* and a novel tumor suppressor gene, IfI44L, which has been implicated in cancer stemness, metastasis, and drug resistance via regulating met/Src signaling (49). Controls include MM marker genes *Msln* (mesothelin) and *Wt1.* Note floxed *Pak2* allele only in *NC;P^-/-^* MMs, with weak wild type (WT) allele due to contaminating stroma in these tumor samples. NMC, normal mesothelial cells.

### Assessment of kinome reprogramming in Pak2-null MM cells using MIB/MS technology

Given that the problem of drug resistance is of paramount importance in aggressive tumors such as MM, we used a technological approach, MIB/MS, that globally measures kinase signaling at the proteomic level (36,37) to assess dynamic reprogramming of the kinome in response to genetic inhibition of Pak2 in MM. This technology was used to compare the kinase expression of early passage MM cells from an *NC;P^wt^* mouse with that of MM cells from an *NC;P^-/-^* mouse, with studies performed in triplicate. Kinome activity, principal component analysis (PCA), identities of up- and down-regulated kinases, and heat-map are depicted in Figure 6A-D. The kinome profiling revealed that as compared to *NC;P^wt^* MM cells, the *NC;P^-/-^* MM cells had multiple kinase changes suggestive of an epithelial to mesenchymal transition (EMT). Prominent among the changes associated with loss of Pak2 were upregulation of Pdgfra, Pdgfrβ, Limk1, Fyn, Jak1, Mkk6, and Slug as well as down regulation of Met and Ddr1. Immunoblot confirmation of expression changes in selected kinases is shown in Figure 6E and Figure 7.

**Figure 6.**
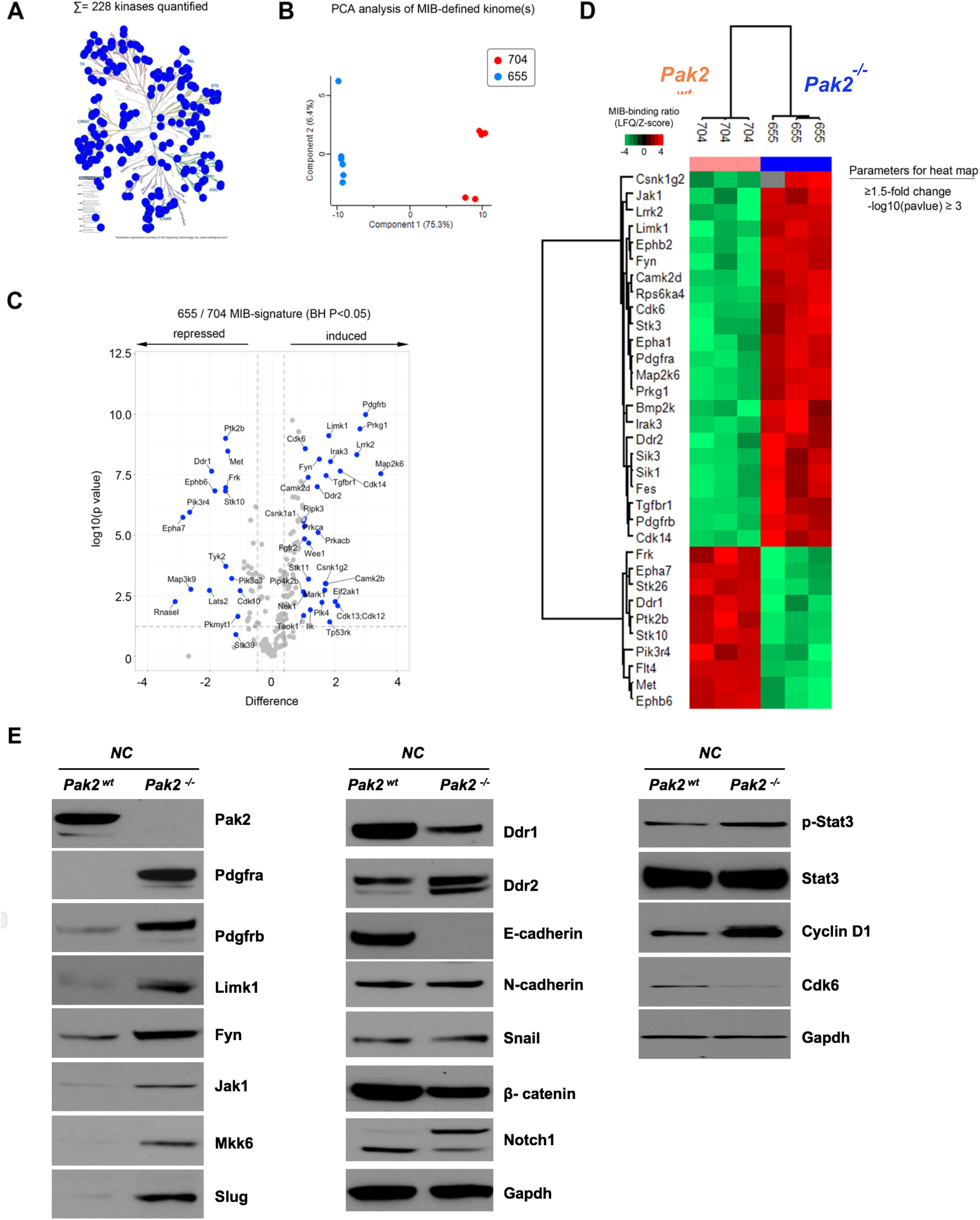
Kinome profiling of *NC;P^-/-^* MM cells and *NC;P^wt^* MM cells. Early passage MM cell cultures were derived from tumors observed in *Nf2^f/f^;Cdkn2a^f/f^* and *Nf2^f/f^;Cdkn2a^f/f^;Pak2^f/f^* mice injected IT with adeno-Cre virus. (**A**) Kinome activity as measure by MIBs. (**B**) PCA analysis, showing differential kinome profiles. (**C**) Volcano plot depicts kinases exhibiting induced or repressed MIB binding in cells with knockout of Pak2. (**D**) Heat-map depicts statistical changes in kinase levels between *NC;P^-/-^* and *NC;P^wt^* MM cells. (**E**) Immunoblot confirmation of expression changes in selected kinases.

**Figure 7.**
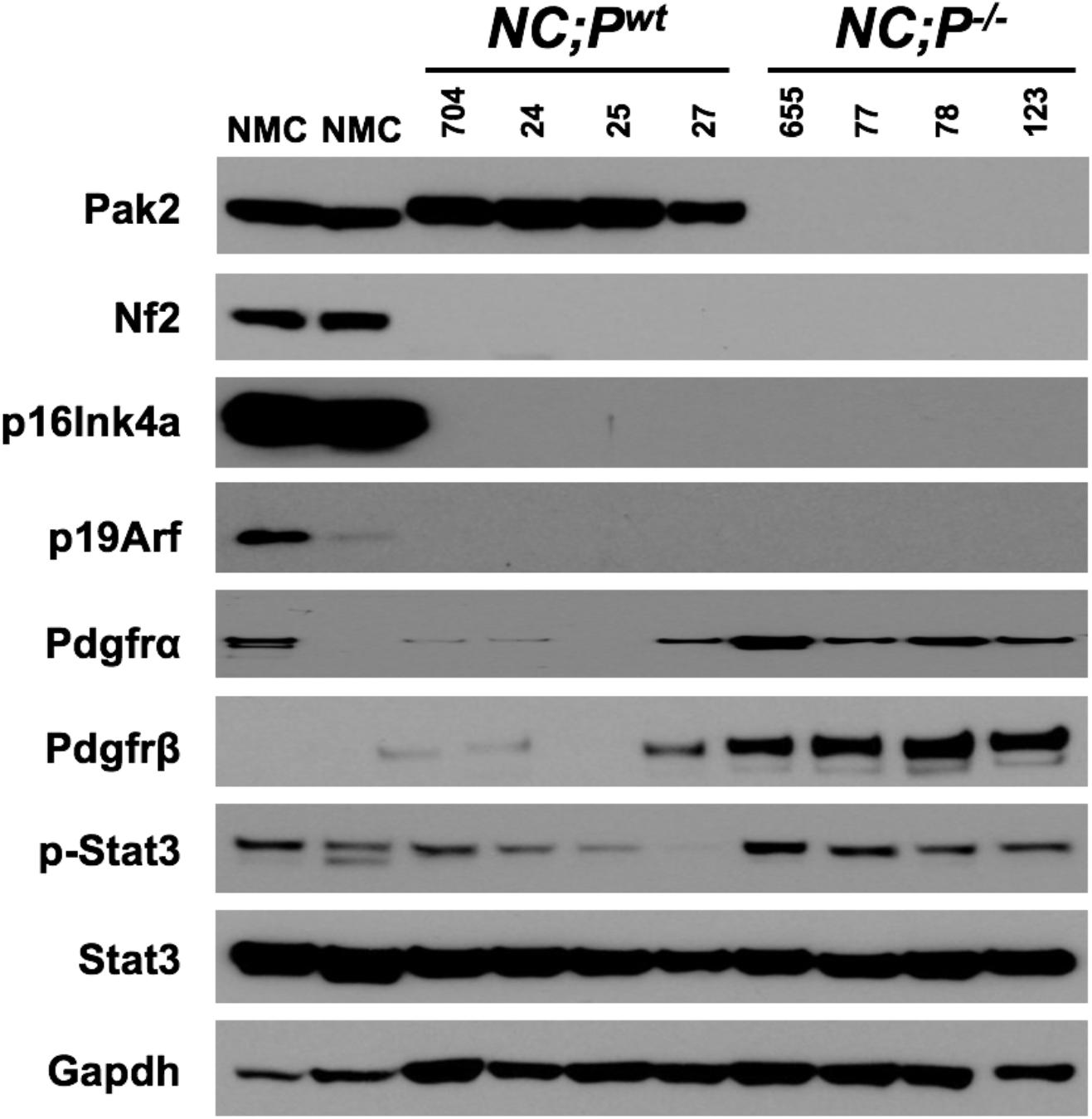
Expression of various key proteins in a series of *NC;P^wt^* and *NC;P^-/-^* cell lines derived from pleural MMs. Loss of Pak2 expression correlates with upregulation of Pdgfra, Pdgfrß, and phospho-Stat3. NMC, normal mesothelial cells.

## Discussion

Losses of *NF2* and *CDKN2A* are thought to play a critical role in MM pathogenesis. In fact, in human peritoneal MM, homozygous deletions in *CDKN2A* and hemizygous loss of *NF2* as detected by fluorescence *in situ* hybridization has been reported to confer a poor clinical outcome, whereas loss of *BAP1* was not associated with clinical outcome (38). In conditional knockout mice, *Nf2^f/f^* mice and *Cdkn2a^f/f^* mice injected IT with adeno-Cre virus resulted in few pleural MMs, whereas *Nf2^f/f^;Cdkn2a^f/f^* mice injected with adeno-Cre virus resulted in aggressive thoracic tumor growth (27,32). In this study, *NC;P^-/-^* mice, with loss of only a single Group I Pak gene, was sufficient to significantly decrease the incidence and delay the onset and progression of both pleural and peritoneal MMs when compared to that of *NC;P^wt^* mice.

Loss of *NF2* results in the direct activation of Group I Paks (20) Group I PAKs are frequently activated in human MM tumors, and genetic or pharmacologic inhibition of PAKs is sufficient to inhibit MM cell proliferation and survival (26). PAK1 is frequently overexpressed in human breast, ovarian, bladder and brain cancers, often secondary to amplification of its 11q13.5-14 chromosomal locus (39). Moreover, breast and bladder carcinoma cells bearing such amplification were shown to be highly sensitive to PAK inhibition by small molecule inhibitors or RNAi, suggesting that these cancer cells are “addicted” to PAK1 overexpression. The role of PAK2 in human neoplasia is less well studied, but it has been linked to a variety of human cancers (23). The pathways that mediate the oncogenic effects of PAK overexpression are not fully known, but include activation of the Erk pathway (via phosphorylation of Raf and Mek), inactivation of *NF2*/Merlin, stabilization of ß-catenin, and possibly by scaffolding interactions that link Pdk1 with Akt (23). Pak1 has been shown to regulate cell motility in mammalian fibroblasts (40), and we show here that loss of Pak2 in *Nf2*-deficient MM cells similarly results in decreased MM cell migration. Loss of PAK function is associated with decreased cell proliferation and migration, with concomitant loss of activity of these signaling pathways (41). Consistent with these data, our findings demonstrate that loss of Pak2 results in diminished MM cell motility, decreased expression of Akt pathway-related genes, and reduced Erk activity.

Our RNA-seq analysis revealed that loss of Pak2 counteracts some aspects of Nf2 loss-of-function in MM cells by downregulating the expression of multiple genes involved in oncogenic pathways, particularly genes encoding proteins involved in Wnt signaling, such as *Wnt9b* and several *Fzd* (frizzled class receptor) genes as well as in Akt signaling, e.g., *Igf2, Fgf, Fgfr,* and *Pik3r3.* Among the comparatively few upregulated genes observed in MM cells from *NC;P^-/-^* mice, *Dkk2* is a Wnt inhibitor gene, *Foxg1* encodes an oncoprotein that inhibits the transcriptional activation of the proapoptotic protein FoxO1 (42), and *Dusp4/5* encode dual-specificity phosphatases that dephosphorylate MAPK proteins such as ERK. Consistent with the latter, we found that Erk activity was markedly diminished in *NC;P^-/-^* MM cells compared to *NC;P^wt^* MM cells (Figure 4B). One other upregulated gene in *Pak2*-null MM cells, *Styk1*, encodes a serine threonine tyrosine kinase 1 that has been correlated with poor prognosis, tumor invasion, and metastasis of non-small cell lung cancer (NSCLC) patients (43). STYK1 overexpression also promoted proliferation, migration, and invasion of NSCLC cells and induced EMT by E-cadherin downregulation and Snail upregulation. Similarly, E-cadherin expression was greatly downregulated in the *NC;P^-/-^* MM cells used for our MIB/MS kinome profiling study, whereas E-cadherin expression was abundant in *NC;P^wt^* MM cells (Fig. 5E). Collectively, these data point to a positive role for group I Paks in tumorigenesis, mediated by several central signaling pathways. Our findings further suggest that inhibitors of Pak might be useful in switching off such signaling pathways. The data presented here help establish a framework for understanding MM adaptation and permit the design of rational combinations of targeted agents, as clinical or near-clinical inhibitors for many protein kinases already exist.

The RNA-seq profiling presented here also demonstrated that numerous genes that are downregulated in *NC;P^-/-^* MM cells are involved in muscle contraction and cardiac epithelial to mesenchymal transition (EMT) pathways. These data are consistent with previous work showing a significant role for Group I Paks in myoblast and cardiac muscle function. For example, the Rac1-Pak2 pathway has been shown to be indispensable for zebrafish heart regeneration (44). Furthermore, Pak2 has been identified as a primary mediator of ER stress in chronic myocardial injury, and melatonin-mediated Pak2 activation has been shown to decrease cardiomyocyte death by repressing hypoxia reoxygenation injury-induced endoplasmic reticulum-related stress (45). In addition, Group I Paks have been shown to promote skeletal myoblast differentiation during postnatal development and regeneration in mice, and adult mice conditionally lacking both *Pak1* and *Pak2* in the skeletal muscle lineage developed an age-related myopathy, with muscles exhibiting centrally-nucleated myofibers, fibrosis, and signs of degeneration (46). As in the RNA-seq studies, kinome profiling assayed by MIB/MS technology also uncovered kinase changes indicative of EMT in *NC;P2^-/-^* MM cells, including upregulation of Pdgfra, Pdgfrb, Fyn, and the EMT transcription regulator Slug. Recent work has clarified the EMT cell plasticity program as a set of dynamic transitional states between the epithelial and mesenchymal phenotypes, with EMT and its intermediary states serving as critical drivers of organ fibrosis and tumor progression (47).

Together, these findings suggest that *Nf2*-deficient MM cells with loss of Pak2 ultimately adapt by reprogramming their kinome to bypass the need for Pak activity. While Paks are vital to oncogenic signaling in *NF2*-null MM cells, targeted genetic inactivation of *Pak2* in *Nf2*-null MM cells resulted in downregulated expression of genes involved in oncogenic pathways. Eventually, however, MMs develop in *NC;P^-/-^* mice apparently by compensatory activation of oncogenic pathways involving other kinases such as Styk1, Pdgfrα/β, Axl, Jak/Stat, and Fyn. Such information can inform the design of rational combination therapies for MM using molecularly targeted inhibitors. Germane to this, one kinase gene *(Axl)* that was upregulated in MMs from *NC;P^-/-^* mice is noteworthy because there are promising new inhibitors for Axl that might be used in combination with a Pak inhibitor (48). The identification of such secondary pathways that could be co-targeted to prevent resistance to anti-Pak drugs, thus sets the stage for future preclinical studies with novel therapeutics.

## Acknowledgments

This work was supported by NCI grants CA148805 (to J.R. Testa and J. Chernoff) and CA06927 (to FCCC) and an appropriation from the Commonwealth of Pennsylvania to FCCC. Other support was provided by the Local #14 Mesothelioma Fund of the International Association of Heat and Frost Insulators and Allied Workers. The following FCCC core services assisted this project: Laboratory Animal, Transgenic Mouse, Genomics, Cell Culture, DNA Sequencing, Histopathology, and Biostatistics and Bioinformatics Facilities.

