## Supplemental Figures 1, 2 and Table 1 for "Inactivation of p21-Activated Kinase 2 (Pak2) Inhibits the Development of *Nf2*-Deficient Malignant Mesothelioma"

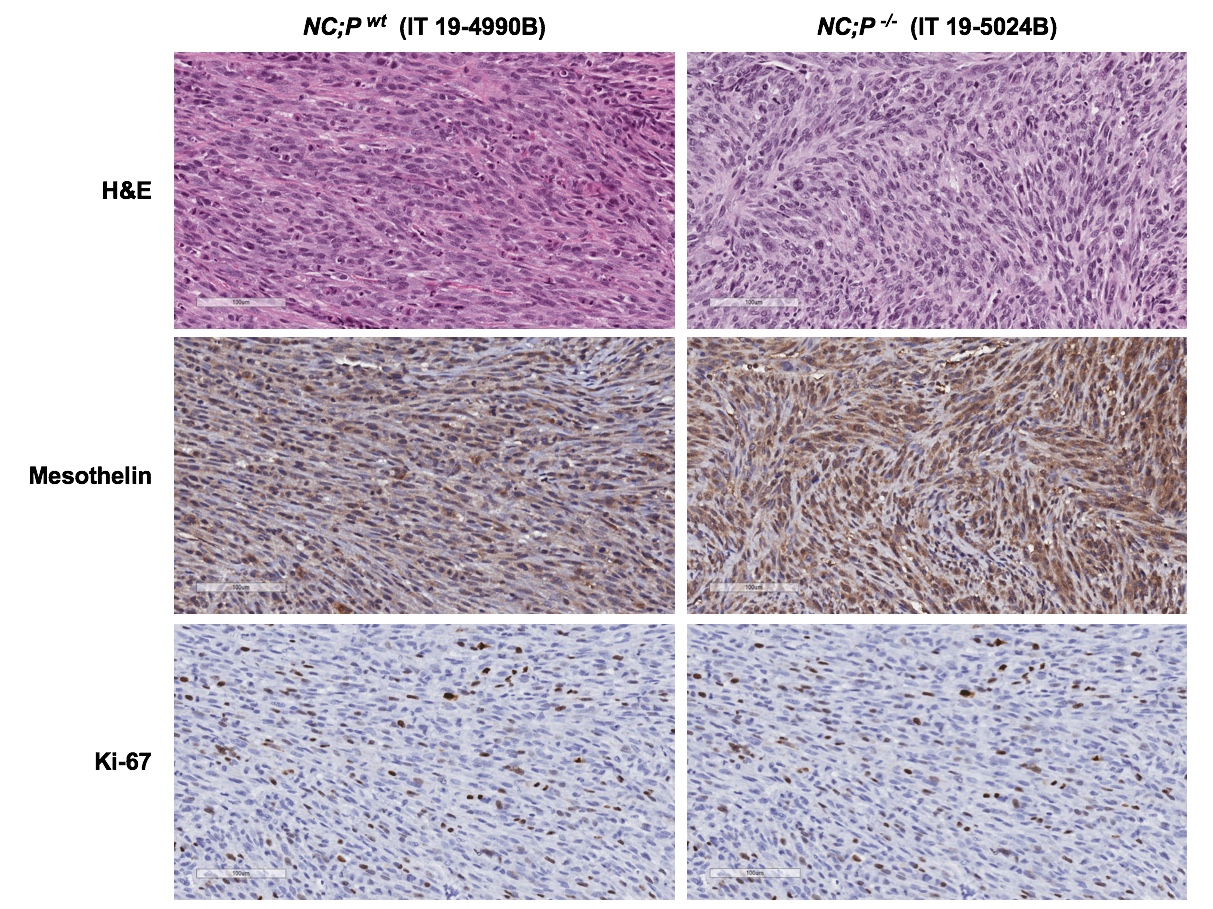

**Supplemental Figure S1.** Intrathoracic MMs from NC;Pak2^wt^ and NC;Pak2^-/-^ mice showed similar histology, with nearly all tumors being sarcomatoid. (**A**) MM from Nf2^f/f^;Cdkna^f/f^;Pak2^+/+^ mouse injected i.p. with adeno-Cre. (**B**) MM from Nf2^f/f^;Cdkna^f/f^;Pak2^f/f^ mouse injected i.p. with adeno-Cre. Bars in left corner of each image represent 100 μm.

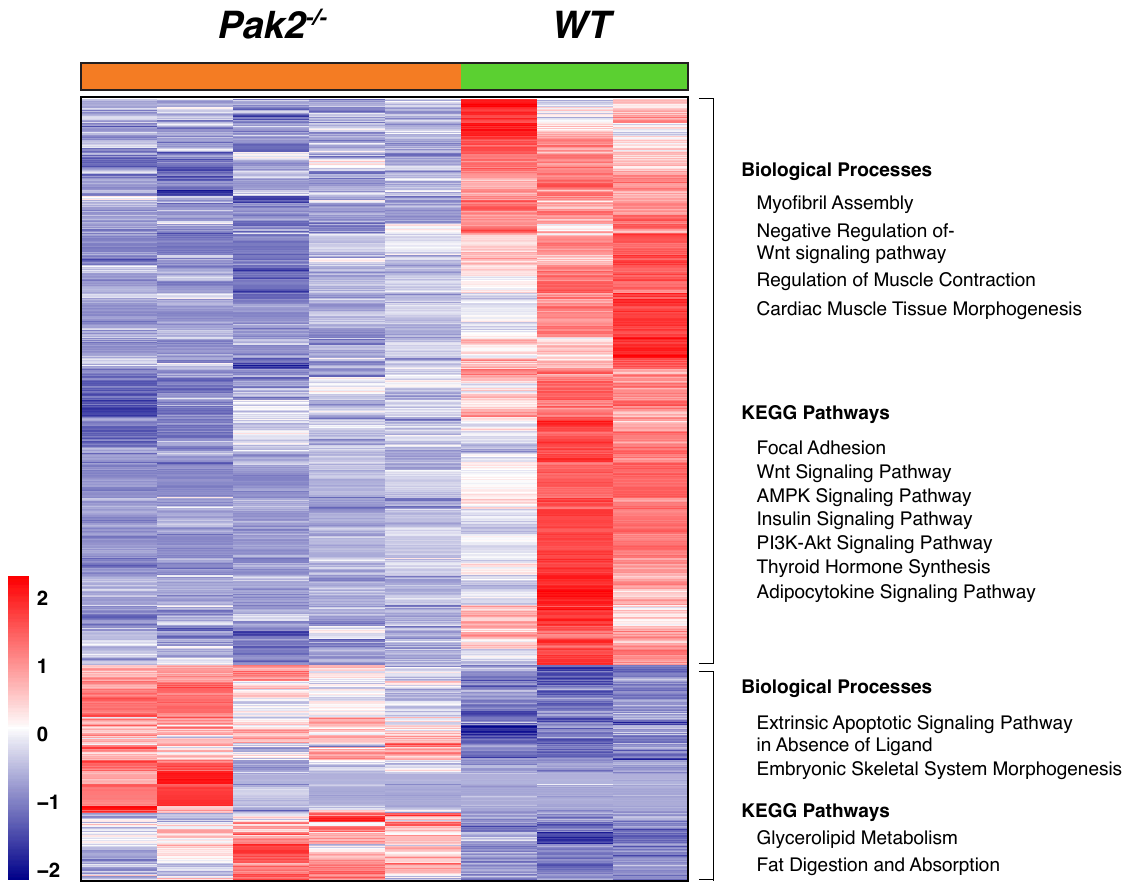

**Supplemental Figure S2.** Heatmap showing list of differentially expressed genes between peritoneal MMs from 3 *NC;P^wt^* mice and 5 peritoneal MMs from *NC;P^-/-^* mice. Using the DESeq2 method(33) to identify differentially expressed genes, 1388 genes were found to be significant (FDR < 5%; Fold-change cutoff log2 >1 or log2 < -1). For these genes, using the mean-centered VST (variance stabilizing transformation), normalized count data are shown as a heatmap. Using DAVID bioinformatics resource,(50) we identified biological processes and KEGG pathways that are enriched for up- and down-regulated genes (p-value < 0.05), as depicted beside the heatmap.

| **Differentially expressed genes identified by RNA-seq analysis of *NC;P^wt^* vs. *NC;P^-/-^* MMs** |  |  |
| --- | --- | --- |
| *Axin2* | Axin-2F | GGCTGCGCTTTGATAAGGTC |
|  | Axin-2R | AGTTCCTCTCAGCAATCGGC |
| *Fgf9* | Fgf9-1F | CTCTGTCTGCAACTGCGGC |
|  | Fgf9-1R | CCGAGGTAGAGTCCACTGTC |
| *Pax7* | Pax7-1F | GAGCATCCTTAGCAACCCGA |
|  | Pax7-1R | TGTGGACAGGCTCACGTTTT |
| *Fzd4* | Fzd4-1F | TGCAGTTCCTCCTGCTCCT |
|  | Fzd4-1R | GCCGATGGGGATGTTGATCT |
| *Fzd9* | Fzd9-1F | GACCGGTTTTGTGGCTCTCT |
|  | Fzd9-1R | CCGCCAGAAGTCCATGTTGA |
| *If144L* | IF144L FOR 3 | 5′-GCTGTGTGATTCAATGGGGC-3′ |
|  | IF144L REV3 | 5′-TAGGACAAAAGCCACGCAGT-3′ |
| *Fgfr4* | Fgfr4 FOR | 5′-AAGGTGGTCAGTGGGAAGTC-3′ |
|  | Fgfr4 FOR | 5′-AGCTGTAGCGAATGCTACCC-3′ |
| *Igf2* | Igf2 FOR | 5′-GTACTTCCGGACGACTTCCC-3′ |
|  | Igf2 REV | 5′-GAGGGAGTGGAGCAGAGAGA-3′ |
| **Mesothelioma marker genes** |  |  |
| *Wt1* | mWT1-1F | CACGGCACAGGGTATGAGAG |
|  | mWT1-1R | GTTGGGGCCACTCCAGATAC |
| *Msln* | mMSLN-2F | CCACACTCCTCCATGCTGTG |
|  | mMSLN-2R | CTGTTGCCCTGGAAACATGG |
| ***Pak2* floxed allele (exon2 deletion)** | mPak2-E1-1F | GTCGGGCGGAGACCTTTC |
|  | mPak2-E3-1R | GCCAGTGAACTCTCCCGTAA |

**Supplemental Table S1.** Primers used for semi-quantitative RT-PCR analysis.
